# *Sox2* is an oncogenic driver of small cell lung cancer

**DOI:** 10.1101/657924

**Authors:** Ellen Voigt, Hannah Wollenzien, Josh Feiner, Ethan Thompson, Madeline Vande Kamp, Michael S. Kareta

## Abstract

Although many cancer prognoses have improved in the past fifty years due to advancements in treatments, there has been little to no improvement in therapies for small cell lung cancer (SCLC) which currently has a five-year survival rate of less than 7%. One promising avenue to improve treatment for SCLC is to understand its underlying genetic alterations that drive its formation and growth. One such mutation in SCLC, which appears in many cancers, is of the *Rb* gene. When mutated, *Rb* causes hyperproliferation and loss of cellular identity. Normally *Rb* promotes differentiation by regulating lineage specific transcription factors including regulation of pluripotency factors such as *Sox2*. However, there is evidence that when certain tissues lose *Rb*, *Sox2* becomes upregulated and promotes oncogenesis. To better understand the relationship between *Rb* and *Sox2* and to uncover new treatments for SCLC we have studied the role of *Sox2* in *Rb* loss initiated tumors by investigating both the tumor initiation in SCLC genetically engineered mouse models, as well as tumor maintenance in SCLC cell lines.

## Introduction

Small cell lung cancer (SCLC) is a devastating disease where survival rates are typically measured in 2-year segments, rather than the typical 5. Its low rate of detection at early stages is exacerbated by its ability for rapid metastasis and almost invariable resistance to therapy (1). Patients who are stricken by this disease face a 6% 2-year survival rate while most will succumb less than a year after diagnosis (2). Despite this alarming statistic, the standard of care for treating SCLC has remained essentially the same for the past 40 years and few innovations have been developed. Patients still rely on platinum-based drugs that are not ideal because of their lack of precision. Just as researchers have found new drugs for many other cancers based on their DNA mutations, scientists have looked for new drugs to treat SCLC. In the pursuit of these new therapies, researchers begin by seeking to understand the underlying genetic causes of SCLC.

As in many other cancers, the retinoblastoma protein (*Rb*) is lost in SCLC (6–12). Normally *Rb* is recognized for its responsibility in the regulation of the cell cycle but it also interacts with many lineage-specific transcription factors (1). One of the transcription factors regulated by *Rb* is *Sox2* (2). Known primarily as a pluripotency factor, *Sox2* is also a key master regulator of neural and neuroendocrine cell types (3–7). As a master regulator, *Sox2* influences cell identity early and widely in the cell fate decisions. Pulmonary neuroendocrine cells are a common cell of origin for SCLC, therefore is possible that *Sox2* activity in neuroendocrine cells following *Rb* loss induces stem or progenitor genetic networks that help to drive oncogenesis. To that end, we generated a conditional knockout mouse in which we could perturb *Sox2* activity in a SCLC mouse model to assess the consequence of SCLC formation after *Sox2* loss. We observed that *Sox2* is indeed required for SCLC formation, validating *Sox2* as an oncogenic driver of SCLC.

## Experimental Methods

### Ethics statement

Mice were maintained according to the guidelines set for the by the NIH and were housed in the Sanford Research Animal Research Center, accredited by AAALAC using protocols reviewed by our local IUACUC.

### SCLC mouse tumor initiation

We modeled SCLC in the *Rb* ^lox/lox^, *p53* ^lox/lox^, *p130* ^lox/lox^, *Rosa* ^luc^ mouse line (8). *Sox2* ^+/+^,^+/lox^, and ^lox/lox^. To study SCLC tumor initiation, we injected Cre-recombinase adenovirus (Baylor Vector Core) into the mouse lungs by intratracheal injection to excise the floxed genes. The mice were assigned to either a six-month cohort, a three-month cohort or the survival curve. Mouse lungs, livers, and any tumors were harvested for immunohistochemistry.

### SCLC cell lines

We used the murine SCLC cell lines KP1 and KP3 and the human SCLC lines NJH29 (H29) and NCI-H82 (H82) (8, 9). The cells were maintained in suspension and cultured in RPMI with 10% bovine growth serum and penicillin/streptomycin.

### SCLC lung and liver immunohistochemistry

The Sanford Research Pathology Core performed the immunohistochemistry for this study. The mouse lungs, livers and tumors were stained with H&E, for Sox2, calcitonin (Sigma, 38198 1:2,000), anti-phospho-histone H3 (EMD Millipore 06-570 1:500), cleaved caspase 3 (CST-9664 1:100), ki67.

### Tissue image analysis and statistics

We used Cell Profiler to count the tumors unbiasedly and to determine whether they are CGRP+ to identify them as SCLC over another kind of cancer.

### Lentiviral and siRNA transfection

We made the lentivirus using the packaging plasmids VSVG, pMDL, and RSV in 293T cells. Virus was concentrated and titired for reproducible transductions. We measure their viability after the knock down with an alamar blue assay, and the levels of apoptosis with Annexin V and flow cytometry. qPCR was used to confirm the knock down of Sox2 in the cells.

## Results

To investigate if Sox2 is required for the formation of SCLS, we bred a mouse line containing a conditional *Sox2* allele (*Sox2*^lox/lox^) to an existing mouse models for SCLC. This mouse, consisting of *Rb*^lox/lox^; *p130*^lox/lox^; *p53*^lox/lox^; *Rosa26*^LSL-Luciferase^ alleles, displays all the hallmarks of human SCLC, mainly the same histological characteristics, rapid metastasis, and chemoresistance (8, 10, 11). To overcome the dramatic effects of *Rb*- and *p53*-loss in the mouse, we localized Cre-mediated recombination to the lung by the intratracheal instillation of a Cre-expressing adenovirus (Adeno-Cre). Due to the presence of the floxed *p130* allele (8), we observed early tumors around 3-months, with a robust tumor burden 6-months after Cre-recombination (Figure 1).

**Figure 1.**
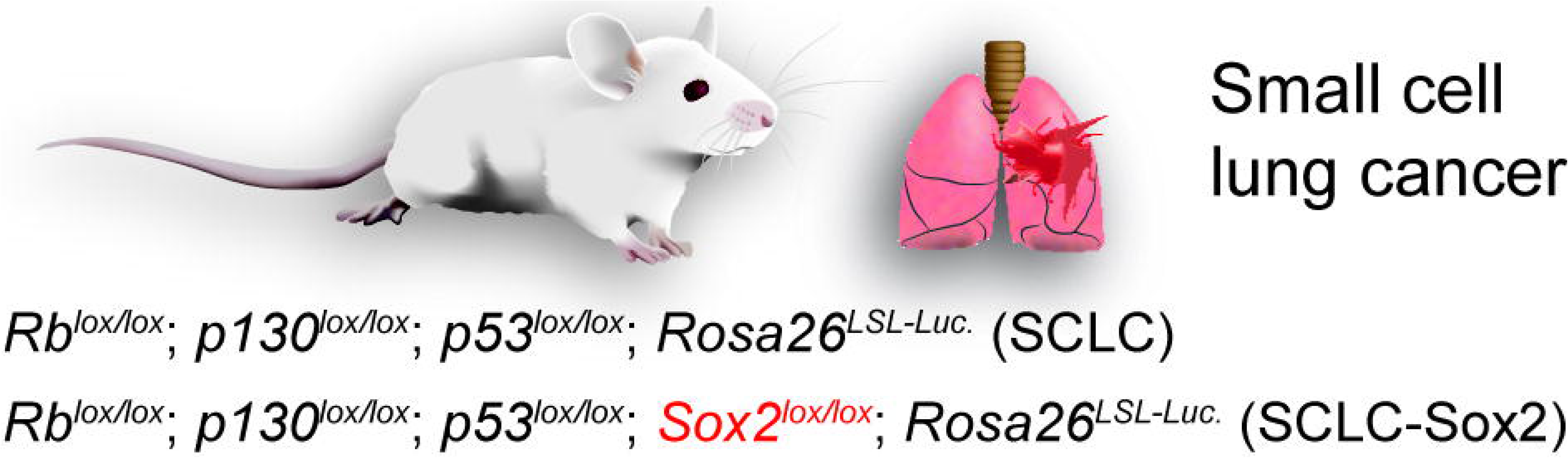
Genetically engineered mouse model for the study of *Sox2* in SCLC.

We bred this SCLC mouse line with a *Sox2*^lox/lox^ line, ensuring that we generated mice of all three potential *Sox2* genotypes: *Sox2*^+/+^, *Sox2*^+/lox^, and *Sox2*^lox/lox^. By this strategy, we can determine if one or both alleles of *Sox2* may be involved in SCLC formation. Tumors were initiated by Adeno-Cre, and as expected *Rb*Δ; *p53*Δ; *p130*Δ mice showed a sizeable number of tumor foci showing the histological characteristics of SCLC 6 months after Cre infection (Figure 2) (8, 12). The *Rb*Δ; *p53*Δ; *p130*Δ; *Sox2*Δ mice had a nearly complete loss of SCLC foci observed at the same timepoint (Figure 2A,B). We observe a significant lengthening of the lifespan of the *Rb*Δ; *p53*Δ; *p130*Δ; *Sox2*Δ mice (Figure 2C). Finally, to fully characterize these tumors and ensure *Sox2* loss in the *Rb*Δ; *p53*Δ; *p130*Δ; *Sox2*Δ mice, we have optimized immunohistochemistry staining and an unbiased image analysis pipeline to identify and characterize these tumors to perform thorough statistical analysis of the *Rb*Δ; *p53*Δ; *p130*Δ; *Sox2*^+^ tumors compared to the few *Rb*Δ; *p53*Δ; *p130*Δ; *Sox2*Δ tumors (Figures 2D and 2E). Indeed, of the few tumors *Rb*Δ; *p53*Δ; *p130*Δ; *Sox2*Δ tumors, a plurality showed immunoreactivity to Sox2 antibodies, indicating that they are the result of incomplete Cre function. However, a small number of *Rb*Δ; *p53*Δ; *p130*Δ; *Sox2*Δ tumors appeared to be *Sox2*Δ indicating that *Sox2* activity may not be necessary in some SCLC tumors.

**Figure 2.**
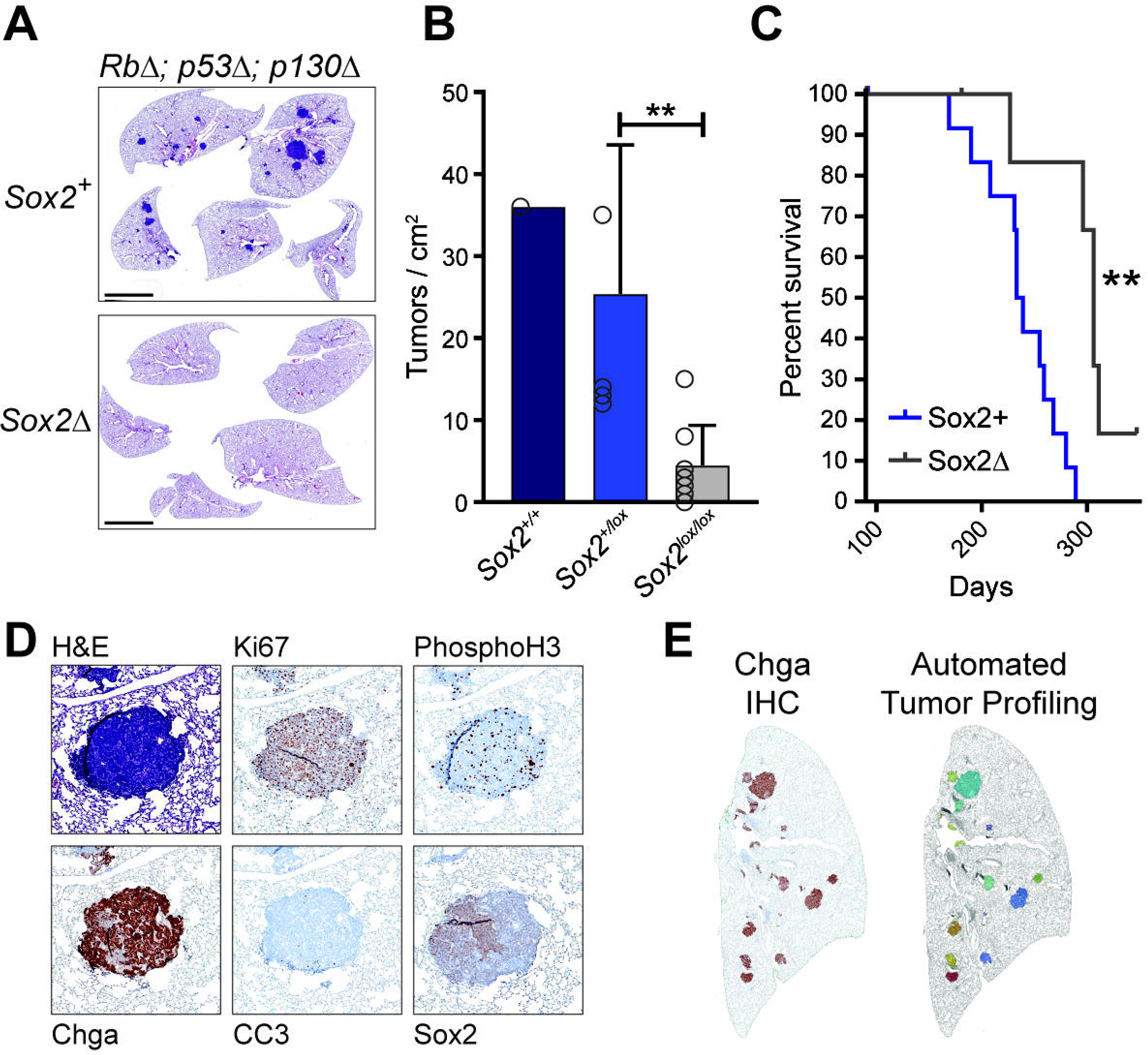
SCLC tumors show a requirement for Sox2. A) SCLC was initiated by intratracheal injection of Adeno-Cre virus and the H&E stained lungs were screened for SCLC foci at 6 months. Scale bar = 4 mm. B) Quantification of tumor foci observed in SCLC mice at 6 months. C) Survival curve of Sox2+ and Sox2Δ mice. D) Immunohistochemistry of tumors for growth markers Ki67, Phospho-histone H3 (PhosphoH3), and cleaved caspase 3 (CC3), and the neuroendocrine markers Chga and Sox2. E) Automated tumor identification by Chga staining, shown as colored areas (right).

To investigate the requirement for the *Rb*-*Sox2* axis in the growth and maintenance of already established tumors, murine cell lines for SCLC were infected with adenovirus harboring *Rb* and a *GFP* reporter (Adeno-Rb-GFP) or *GFP* alone (Adeno-GFP). After sorting for GFP^+^ cells, the SCLC cell lines showed reduction of *Sox2* upon overexpression of *Rb* (Figure 3A). As *Rb* overexpression in mouse SCLC will slow the growth of established tumors (13), we sought to determine if knockdown of *Sox2* would have deleterious effects on the cancer cells. Indeed, we observed that short hairpin-mediated knockdown of *Sox2* (*shSox2*) achieve a ∼60-90% knockdown by RT-qPCR (Figure 3B) and resulted in an increase in cell death (Figure 3C). *Sox2* knockdown also dramatically reduced the growth of both mouse and human SCLC cell lines (Figure 3D).

**Figure 3.**
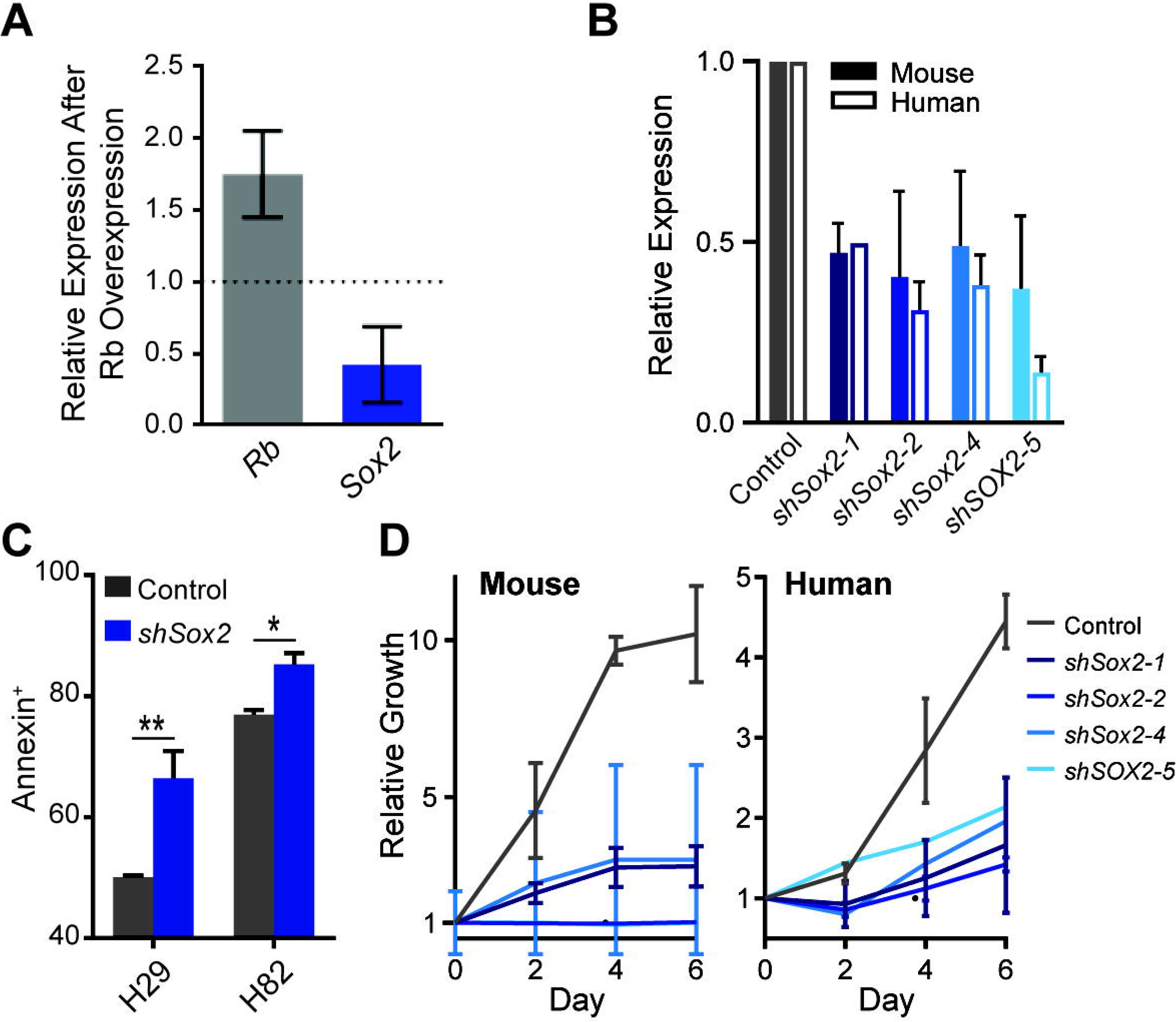
Rb regulates Sox2 in SCLC and affects cellular viability and growth. A) Expression of Rb and Sox2 in GFP+ sorted SCLC cell lines infected with Adeno-Rb-GFP relative to Adeno-GFP. B) Knockdown efficiency of 3 murine Sox2 shRNAs (shSox2-1,2,4) and one human hairpin (shSOX2-5), plotted relative to a non-targeting hairpin control. C) Annexin V labeling of apoptotic cells after transduction with shSox2 or control hairpins in two human SCLC cell lines. High numbers of apoptotic cells are due to residual dead cells after Puromycin treatment to select transduced cells (shSox2 or controls). Due to the non-adherent nature of these cell lines, dead cells are difficult to fully remove before further testing. D) Alamar blue assay to measure growth of mouse and human SCLC cell lines after transduction with Sox2 hairpins or controls. Error is the biological variance between two different murine (KP1 & KP3) or human cell lines (H29 & H82).

## Discussion

*Sox2* is an essential transcription factor, and has been observed to be misregulated in various cancer of the epithelium. As a cancer that rises from the lung epithelium, it seemed reasonable that *Sox2* may indeed be a driver of SCLC (12). Furthermore, we had previously observed that Rb specifically binds to and repressed the core pluripotency network, including *Sox2* (2). As SCLC is initiated in part, by loss of *Rb* it was possible that *Rb*-loss resulted in *Sox2* upregulation, which might drive SCLC (10, 14, 15). Indeed, SCLC oftentimes shows amplification of *Sox2* (14). Therefore, this data and others shows that *Sox2* activity is indeed a driver of SCLC (14). We did, however, observe a small minority of *Rb*Δ; *p53*Δ; *p130*Δ; *Sox2*Δ tumors. Whether these tumors were indeed independent of Sox2 activity, or if they received signaling from the microenvironment which activated *Sox2* target genes in the absence of *Sox2* is currently unclear.

One potential interesting facet of SCLC biology is the realization that it is not a homogenous disease; rather it is characterized by four different subtypes (16). The *Rb*^lox/lox^; *p130*^lox/lox^; *p53*^lox/lox^ mouse model predominantly gives rise to the most prevalent subtype of SCLC, SCLC-A (17). SCLC-A is characterized by strong expression of the neural bHLH transcription factor, *Ascl1* (18, 19). In neural cells, *Ascl1* is a direct target of *Sox2* (20–22). Therefore, in SCLC, a neuroendocrine tumor, *Sox2* activity may upregulate *Ascl1*. *Sox2* may therefore may bridge the loss of *Rb* and *Ascl1*. However, the exact mechanism that *Sox2* may have in the regulation of *Ascl1*, or in the major genetic determinants of the other SCLC subtypes is currently unclear. What these results do indicate is that *Sox2* is required for SCLC, and may in the future may be a potential biomarker or therapeutic target towards the treatment of this devastating disease.

## Acknowledgements

We would like to acknowledge the NIGMS Center for Pediatric Research for funding support (5P20GM103620-04). Also the Sanford Research Pathology and Flow Cytomotry Cores, which are supported by the NIGMS (P20GM103548).

